# Contingent-behavior assay to study the neurogenetics of addiction shows zebrafish preference for alcohol is biphasic

**DOI:** 10.1101/2021.05.04.442404

**Authors:** Fatima Megala Nathan, Caroline Kibat, Tanisha Goel, James Stewart, Adam Claridge-Chang, Ajay S. Mathuru

## Abstract

Alcohol use disorders are complex, multifactorial phenomena with a large footprint within the global burden of diseases. Here, we report the development of an accessible, two-choice self-administration zebrafish assay (SAZA) to study the neurobiology of addiction. Using this assay, we first demonstrated that, while zebrafish avoid higher concentrations of alcohol, they are attracted to low concentrations. Pre-exposure to alcohol did not change this relative preference, but acute exposure to an alcohol deterrent approved for human use decreased alcohol self-administration. A pigment mutant used in whole-brain imaging studies displayed a similar relative alcohol preference profile, however, mutants in *CCSER1*, a gene associated with alcohol dependence in human genetic studies, showed a reversal in relative preference. The presence of a biphasic response (hormesis) in zebrafish validated a key aspect of vertebrate responses to alcohol. SAZA adds a new dimension for discovering novel alcohol deterrents, and studying the neurogenetics of addiction using the zebrafish.

## Introduction

Substance dependence is a leading preventable cause of economic loss and premature death [1,2]. One of the most widely abused, legally available psychoactive agents is alcohol [3,4]. In 2019 alone, the consumption of alcohol was estimated to have contributed either directly or indirectly towards 3 million deaths (WHO, 2019). The complex interaction of genetics, the environment, and culture contributed to this high number of preventable deaths and societal loss [5]. The underlying neurobiological mechanisms that result in alcohol use disorders (AUD) are an area of active investigation and involve many neurochemical and neuromodulatory systems [6,7]. Numerous efforts to develop appropriate animal models, in which the cellular, neurobiological, and physiological aspects of alcohol dependence can be studied, are also ongoing [8]. Contingent behavioral assays in non-human animals are considered essential to appropriately model human AUD [9]. Most studies currently use rodents and monkeys for such experiments. These can be limiting because of the costs associated with experimentation as well as the type, and the ease with which certain manipulations can be performed.

The zebrafish, a vertebrate with approximately 10^7^ neurons in the larval stages, is a useful alternative that is being used extensively to dissect the cellular, genetic, and molecular bases of complex brain disorders [10]. Along with their benefits in facilitating *in vivo* drug screens, genetic manipulation, and the interrogation of neural circuits at the level of the whole brain [11–13], zebrafish are also suitable subjects for studying the neurobiology of substance dependence. Most subcortical circuits and brain regions implicated in addiction in mammals are conserved at the level of gene expression in the zebrafish [14], and behavioral effects relevant to a vast array of drugs of abuse [15–17], including alcohol [18–20], have been described. Chronic exposure models have also been developed that examine long-term neuroadaptations and neuroplasticity associated with drug use [21,22]. A majority of substance abuse studies in zebrafish, however, have used non-contingent assay designs [15,23], and only a few have adopted experimental designs that test the choice of the animal directly [24,25]. A paucity of assays where complex behavioral procedures can be implemented to study different phases of development of dependence, such as the contingent presentation of a drug, has been described as a major limitation of this system to model addiction [9]. In general, compared to a non-contingent assay that measures the effects after exposure to the drug, behavioral studies of operant response or active participation are considered to have a better construct validity to model human drug dependence [9,26–29].

Here, we report the development of a new method of active administration for juvenile zebrafish, the Self-Administration Zebrafish Assay (SAZA). Unlike previous designs for adults, we intended to develop SAZA for younger fish, closer to the age range at which *in vivo* neural activity imaging of the whole brain is conducted [11–13]. SAZA at this age also facilitates rapid neurogenetic analyses in studies using CRISPANTS [30,31]. Employing this assay, we found that young zebrafish showed a biphasic response, i.e., an inverted-U-shaped preference, to increasing concentrations of alcohol. The prevalence of such a dose-response curve to addictive substances previously reported only in mammals and birds [32] is thus extended to zebrafish. The sensitivity and the utility of this assay system to conduct neurogenetic studies became apparent when the responses to two genetic mutant lines were examined. Among these, the mutants generated in the Coiled-Coil Serine Rich Protein (*CCSER1;* HUGO gene ID HGNC:29349) gene for this study also demonstrated a way to evaluate the potential pathogenicity of candidate genes discovered in human genetic association studies rapidly using an animal model.

## Results

### A Two-choice self-administration zebrafish assay (SAZA)

We designed a new assay system that contained stimulus delivery zones at one end, a constant influx of fresh system water at the opposite end, and extraction from the middle (Figure 1A). This arrangement created three chemically separable virtual zones that permitted unhindered physical swimming access to the juvenile fish (Figure 1A, Supplementary Video 1). For the experiments described here, our custom-written software in LABVIEW called CRITTA [33] was configured to create a closed-loop system, such that the entry of the fish into the virtual delivery zones triggered a pinch valve system (Figure 1B). Subjects were used only once, and each trial lasted 24 minutes (Figure 1C) divided into three phases – pre-exposure, self-administration, and post-exposure. To eliminate bias due to the stimulus delivery location, the stimulus delivery was randomly assigned to the left for half of the subject fish in a group, and to the right for the other half. A food dye used as a stimulus showed that upon each entry into the stimulus zone fish were briefly exposed to a 5-10 fold diluted stimulus (Figure 1D, E; see methods for details). The volume of the stimulus delivered was proportional to the time fish spent in the stimulus zone (Figure 1F). Even though the dispersal of the stimulus in the tank was dynamic, dependent on the locomotion behavior of each subject fish, it was usually rapidly diluted to a fraction of the concentration detectable inside the delivery zone (Figure 1E). Thus, fish could freely choose to self-administer either a stimulus, or a control solution.

**Figure 1.**
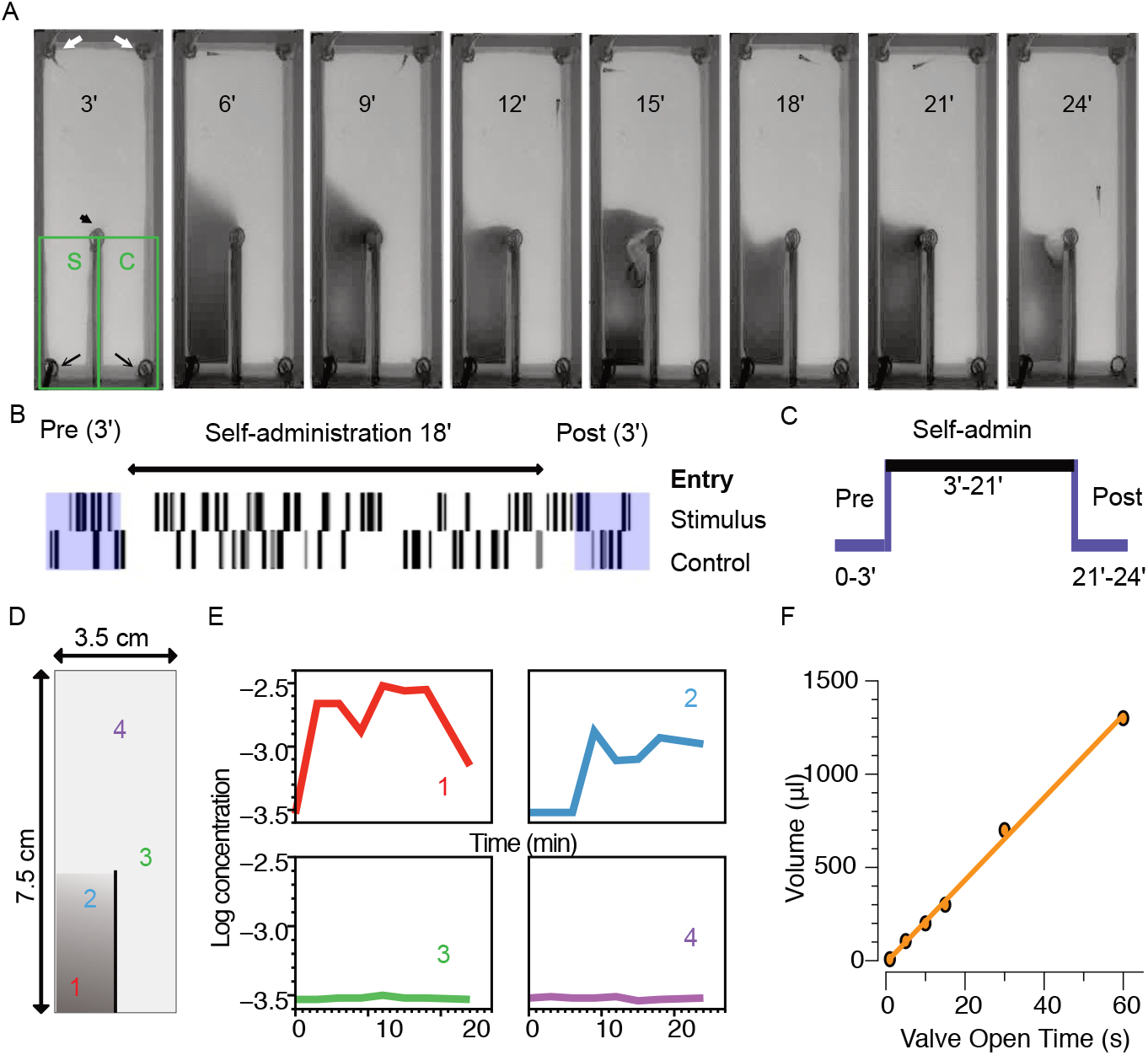
A Two-Choice Self-Administration Zebrafish Assay (SAZA) (A) 3- to 4-week old zebrafish swam freely in the SAZA arena (shown in D) and triggered the delivery of stimulus or system water from tubes (black arrows) in the delivery zones. Fresh system water was delivered at a constant rate of 1 ml/min from one end (white arrows), and water was extracted in the middle (arrowhead). Snapshots show a representative experiment using stimuli containing food dye. Green boxes show stimulus (S), or control (C) delivery virtual zones (B) Sparklines plot for the representative experiment shows the entry of the test subject into the stimulus (top) or control (bottom) zones. The thickness of the sparklines indicates the duration the individual remained in the zone. (C) Stimulus delivery was only triggered during the self-administration phase (self-admin) for 18 minutes in each trial. (D) Water samples from four regions of the arena, which were collected every 3 minutes to estimate the spread of the stimulus, showed (E) a limited spread outside the stimulus zone and a 5-10 fold dilution of the colored stimulus within the stimulus zone during the self-administration phase. (F) The volume of stimulus or control delivered by the delivery tubes was linearly proportional to the duration the fish remained in the delivery zone. Volumes delivered during 1, 5, 10, 15, 30, and 60 seconds of the valve being open are plotted.

### Juvenile zebrafish show a hormetic response to alcohol

A five-point dose-response curve with 0% to 70% alcohol (v/v) as the stimulus was constructed to examine the response of juvenile zebrafish in the SAZA system. For each alcohol dose, 28 fish naive to the SAZA and to alcohol were used. On average, subject fish exposed themselves to the stimulus only for brief periods of a few seconds (Supplemental Figure S1). The relative preference, calculated as the preference index (PI; see methods), was used as the primary measurement such that a PI of + 1 indicated maximum preference for the stimulus, while a PI of −1 indicated maximum avoidance of the stimulus. As expected, the group average PI was approximately equal to 0 when system water was used as a stimulus (Figure 2A, 0% stimulus, blue circles). In this case, on average fish spent 21.6% of the total time in the stimulus zone during the stimulus delivery phase (Figure 2B; 95% CI [27.63%, 15.67%]). In comparison to administration of 0% alcohol, both the PI (Figure 2A; mean difference = 0.3, 95% CI [0.1, 0.4], *p* < 0.004) and the time spent in the stimulus zone (Figure 2B; mean difference = 16.75%, 95% CI [9.34%, 25.10%],*p* < 0.001) increased significantly when the fish administered 5% alcohol. On the other hand, these values showed a significant decrease when the fish had a choice of administering 10% or higher concentrations of alcohol (Figure 2B). The fish showed the greatest avoidance of 70% alcohol: the PI (Figure 2A; mean difference = −0.7, 95% CI [−0.8, −0.4], *p* < 0.001) and the time in the stimulus zone (Figure 2B; mean difference = −8.32%, 95% CI [−12.8%, −3.9%]), *p* < 0.0001) both showed a sharp decrease. That is, fish actively avoided 70% alcohol by choosing to administer the control or the system water. The mean difference between the shared control of 0% stimulus and each of the five stimulus conditions, and the statistical analyses, are tabulated in Supplementary Table 1.

**Figure 2.**
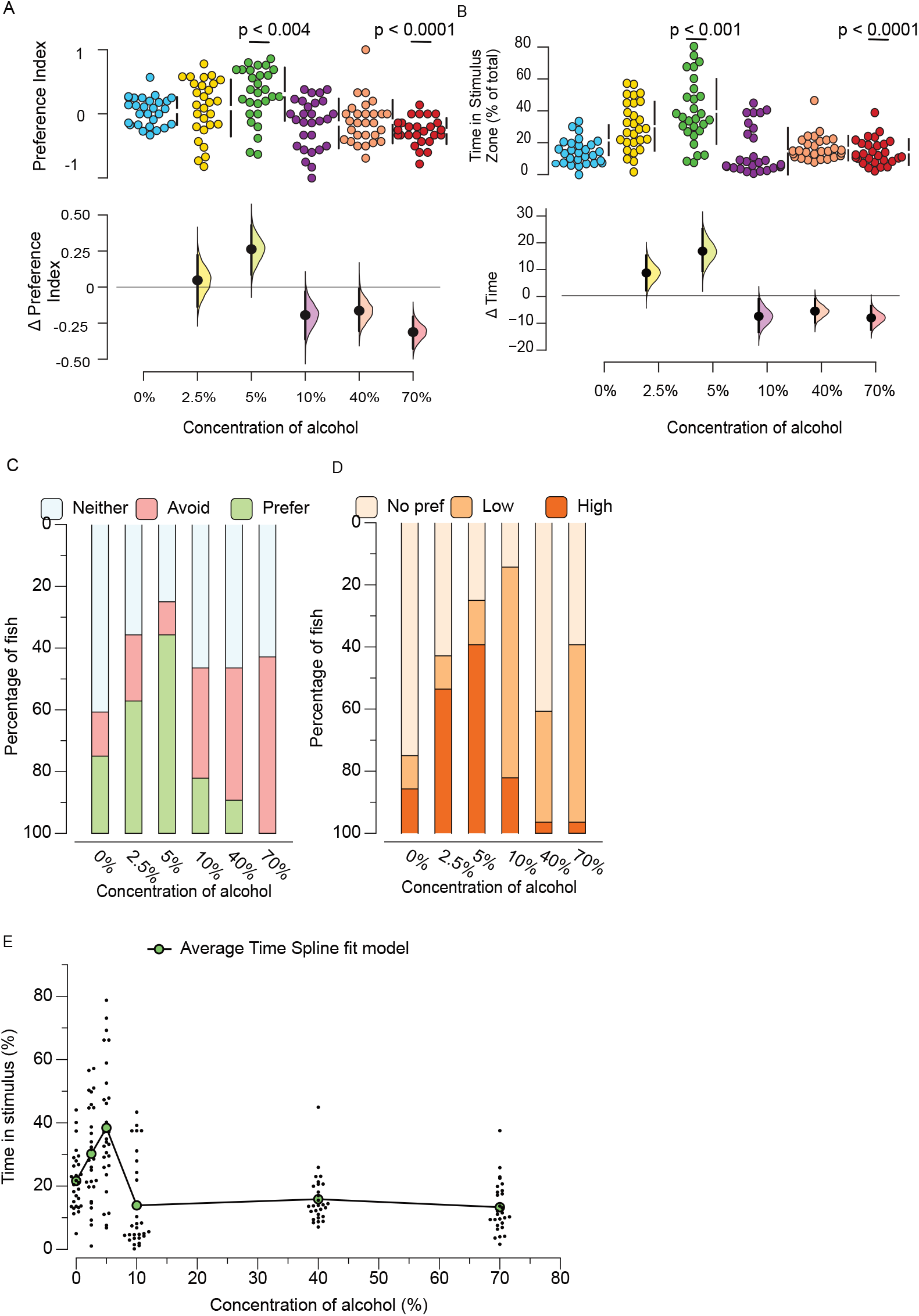
Dose-response to alcohol administration. (A) Preference Index (PI) and (B) percentage time in stimulus zone for five concentrations of alcohol. Top panels show distribution (n = 28/group), and the mean and standard error of the mean are represented adjacent to each group as the gap and black bars, respectively. The bottom panels show the mean difference between different concentrations and the shared control (0%). The mean difference is depicted as a black dot, and the 95% confidence intervals are represented by vertical error bars. The shaded area shows the bootstrap sampling distribution of the mean difference. Stacked bar charts of the percentage of fish in each group showing when they (C) preferred (PI > +0.2) or avoided (PI < −0.2), or (D) spent long (>31.1% of total time) or short (<13.6% of total time) amounts of time in the stimulus zone. (E) Spline fit of average time spent in the stimulus zone shows an inverted-U, or an inverted-J-shaped response, first increasing then decreasing with increasing concentration of alcohol. *p*-values for the permutation t-tests comparing each condition with shared control 0% are reported in Supplementary Table 1. *p*-values for 5% and 70% alcohol that show the largest effects are depicted in (A) and (B).

Based on the responses to the stimulus condition of 0% alcohol, we categorized a PI greater than 1 standard deviation (SD = 0.2) from the mean as a display of clear preference and that of less than 1 SD as avoidance. In-between values were considered to represent no choice. We plotted the proportion of fish showing each of these three preferences (Figure 2C), which revealed that the proportion of fish that showed a preference for alcohol compared to water was biphasic, first increasing and then decreasing with increasing concentrations of alcohol. The same result was seen if the fish were categorized instead according to the amount of time spent in the stimulus zone. Once again, based on their response to 0% alcohol, we categorized the fish that spent 1 SD longer than the mean time in the stimulus zone as high, 1 SD less as low, and the rest as neutral (Figure 2D). A spline fitting model further suggested that juvenile zebrafish showed a biphasic response to alcohol self-administration, or showed hormesis (Figure 2E). This result is in agreement with observations of other biphasic behavioral changes (Supplementary Video 2), such as shoaling [34] and locomotion [19] to increasing concentrations of alcohol. It is also in agreement with concentration-dependent changes in alcohol self-administration reported in other vertebrates [35].

### Zebrafish inbred lines show a similar relative preference for low concentrations of alcohol

The PI for the *nacre^-/-^* line showed a marked reversal, from a positive value at 5% alcohol to a negative value at 10% alcohol, with a very large effect (Figure 3A; mean difference in PI = −0.4, 95% CI [−0.6, −0.2], *p* < 0.001); therefore, we used these two stimuli concentrations for subsequent experiments. The number of entries into the alcohol stimulus zone, the total time spent there, and the average time per entry also decreased at the higher concentration, as shown by the Forest plot in Figure 3B (see, Supplementary Figure S1 for full data).

**Figure 3.**
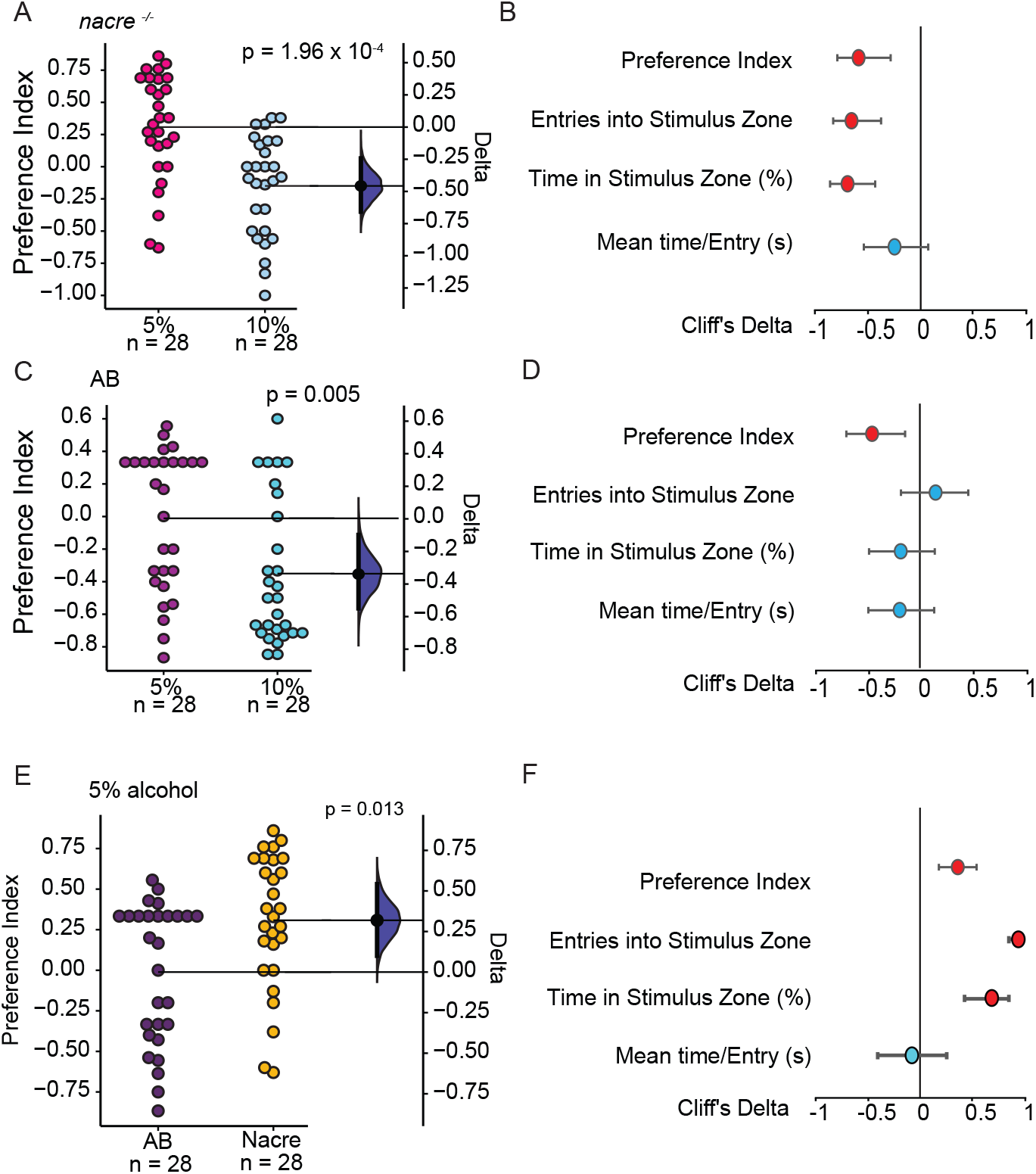
Zebrafish are attracted to low concentrations of alcohol. Both (A, B) *nacre^-/-^* and (C, D) AB WT inbred lines showed a preference for 5% alcohol and avoided 10% alcohol. The mean difference is depicted as a black dot, and the 95% confidence intervals are shown as vertical error bars. The shaded area shows the bootstrap sampling distribution of the mean difference. *nacre^-/-^* Forest plots for (B) *nacre^-/-^* and (C) AB WT show Cliff’s delta (*δ*) comparing 5% alcohol and 10% for other parameters, which are also shown in Supplementary Figure 1 as full data. (E) *nacre^-/-^* showed higher preference for 5% alcohol than AB WT. (F) Forest plots show Cliff’s delta (*δ*) that compared AB WT with *nacre^-/-^* for other parameters. Meaningful effects in Forest plots (−0.4 > *δ* > 0.4, with *p*-values < 0.01) are shown in red.

These experiments were performed in a zebrafish line homozygous for *nacre^-/-^*, as they are used in neural activity imaging studies in many labs, including ours [36–38]. To evaluate if the findings of biphasic alcohol preference are generalizable, we next examined the commonly used AB wild-type fish. AB wild-type fish also showed a similar decrease in preference to administering 10% alcohol compared with 5% alcohol (Figure 3C, mean PI difference = −0.3, 95% CI [−0.5, −0.1], *p* = 0.005). However, unlike in the *nacre^-/-^* line, the other parameter changes showed only minor effects (Figure 3D). We compared the AB and *nacre^-/-^* lines directly to see if they had quantifiable differences (Supplementary Figure S2), and this analysis revealed that the absolute volume and PI for administering 5% alcohol were indeed lower for the AB wild-type fish (Figure 3E, 3F). This result agrees with other behavioral differences reported between the two inbred lines [39]. These strain differences notwithstanding, in general, juvenile zebrafish of both inbred lines showed a relative preference for a brief exposure to 5% alcohol but avoided exposure to 10% alcohol.

### Pre-exposure to alcohol does not increase preference for alcohol self-exposure

Previous studies have reported an increase in conditioned place preference (CPP) in zebrafish adults after a single episode of alcohol exposure at a particular location [40]. This has been interpreted sometimes as reflecting alcohol’s reinforcing effects in fish [40,41]. However, as suggested recently, only contingent procedures are uniquely capable of measuring reinforcing effects [9]. Therefore, to test if pre-exposure to alcohol increased preference for it, in the third experiment, we examined the responses of fish in the SAZA after a multi-day exposure to a low concentration of alcohol.

To evaluate the duration of exposure and the concentration of alcohol needed for juvenile fish to show an effect, pilot experiments were conducted using a range of concentrations (0.3-1% alcohol). Increased shoaling was seen within 5 minutes at lower concentrations, resembling the description of increased shoaling after longer exposure [34]. Exposure to 1% alcohol resulted in a biphasic response (Supplementary Video 2), with clear and pronounced effects within minutes. Fish first showed increased social cohesion, followed by episodes of disoriented swimming, swimming ventral up (upside-down) or side-ways, and reduced social cohesion. Fish rapidly recovered normal swimming behaviors once returned to the home tanks without alcohol.

Based on these results, batches of 28 juvenile fish of each strain were pre-exposed daily to 1% alcohol or system water for 30 minutes for 5 days and were tested using the SAZA on the 8th day. We examined their responses to both 5% and 10% alcohol and compared them to those of control fish exposed only to the system water. We performed this experiment for both the inbred lines. Despite our expectations, the responses of the pre-exposed and unexposed fish were comparable (Figure 4). Pre-exposure did not change the PI of the *nacre^-/-^* for 5% alcohol (Figure 4A) and led to only a marginal increase in the preference for 10% alcohol (Figure 4B). A decrease in the time spent in the stimulus zone when 5% alcohol was available (Figure 4C) was accompanied by fewer visits to the control zone (data not shown) and, thus, overall preference was unchanged. In a similar manner, the pre-exposed AB fish also did not show a notable increase in preference for either 5% (Figure 4E) or 10% alcohol (Figure 4F). The only consistent difference was an increase in the frequency of visits to the stimulus zone (Figure 4G, 4H). This suggests that pre-exposure had a minor effect, but this did not translate to an increase in overall tendency to administer more alcohol.

**Figure 4.**
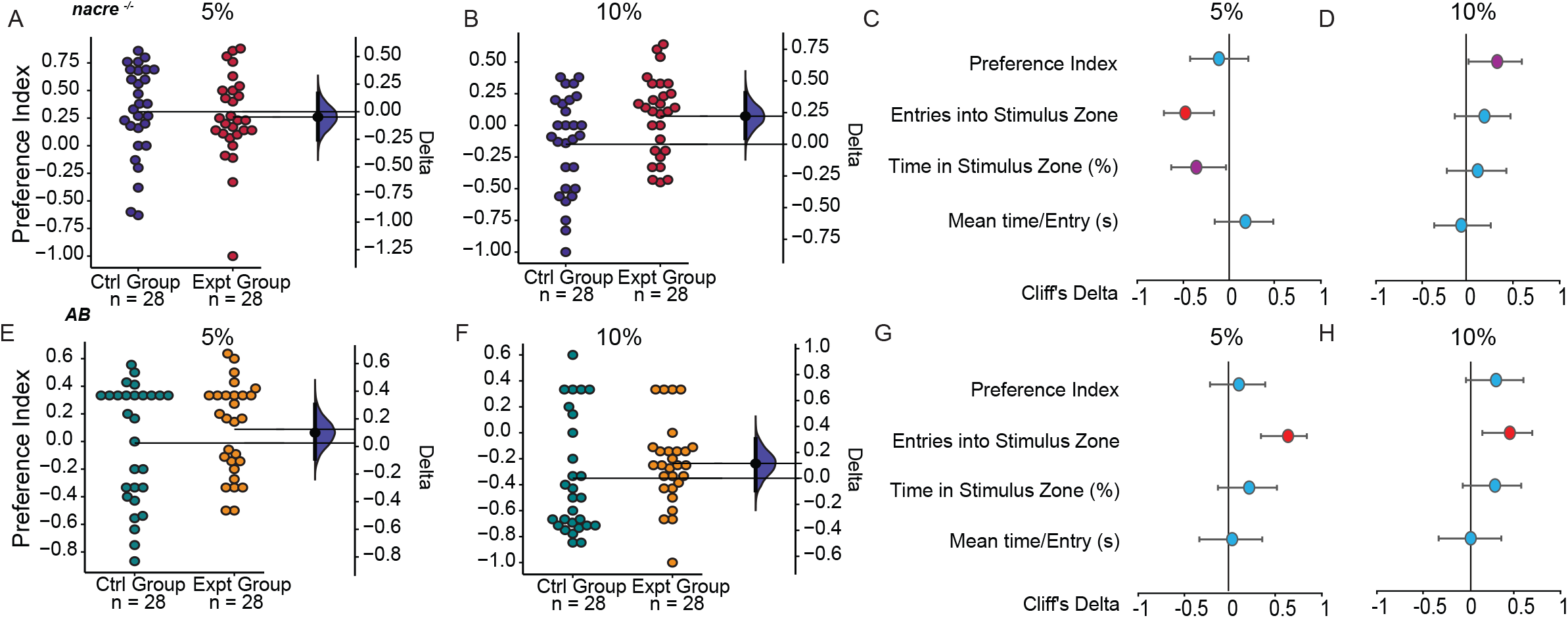
Preference for alcohol does not change with pre-exposure. Both *nacre^-/-^* E, F) AB WT (E, F) strains showed a similar preference for (A) and (C) 5% alcohol and avoidance of (B) and (F) 10% alcohol. The mean difference is depicted as a black dot, and the 95% confidence intervals are shown as vertical error bars. The shaded area shows the bootstrap sampling distribution of the mean difference. Forest plots for nacre-/- (C) and (D) and for AB WT (E) and (F) show Cliff’s delta (*δ*) for other parameters. Meaningful effects (−0.4 > *δ* > 0.4, with p-values < 0.01) are shown in red.

We additionally analyzed if the relative preference for the two concentrations changed after pre-exposure to alcohol. Both *nacre^-/-^* and AB wild-type fish (Supplementary Figure S3A, S3C) continued to prefer 5% over 10% alcohol. The original difference in higher preference for 5% alcohol became more pronounced for the AB wild-type fish (Supplementary Figure S3C, S3D). Together, these observations suggest that pre-exposure to alcohol does not reduce the avoidance of 10% alcohol. Furthermore, the response of AB wild-type fish and *nacre^-/-^* line were qualitatively similar.

### SAZA as a behavioral screen for deterrents and cessation molecules

Disulfiram, or Antabuse, is one of three FDA-approved drugs used as pharmacological deterrents to alcoholism [42]. One proposed mechanism of action is the increase in acetaldehyde, especially in the liver, due to the drug interfering with the breakdown of alcohol [43,44]. The resulting tachycardia, palpitations, headache, panic, and anxiety, among other negative somatic responses, are expected to induce the effect of deterrence. However, recent studies suggest other sites of action of this drug are include changes to dopamine metabolism in the central nervous system [45]. Furthermore, overnight exposure to disulfiram has been shown to inhibit alcohol-induced acute locomotion changes in the zebrafish [46]. In the fourth experiment, therefore, we examined if the SAZA system can be used as a screening tool for drugs that can act as deterrents. As fish prefer to self-administer 5% alcohol, this concentration was used to examine their responses, while control fish were exposed to a vehicle in the form of system water. Disulfiram-treated AB fish spent less time in the stimulus zone (Figure 5A) and made fewer entries into the stimulus zone (Figure 5B). Their PI also only showed a moderate reduction (Figure 5C), and the smaller change in PI was a consequence of the reduced frequency of entry into both the stimulus and control delivery zones (Figure 5D). The *nacre^-/-^* fish phenocopied this effect to some extent. The total stimulus exposure time (Figure 5E) and the number of entries did not change significantly; however, the overall preference for alcohol decreased (Figure 5F). Therefore, disulfiram treatment resulted in a moderate to a large decrease in preference for the normally preferred 5% alcohol for both lines. This suggests the SAZA is a reliable tool for screening novel compounds that can be used as alcohol deterrents.

**Figure 5.**
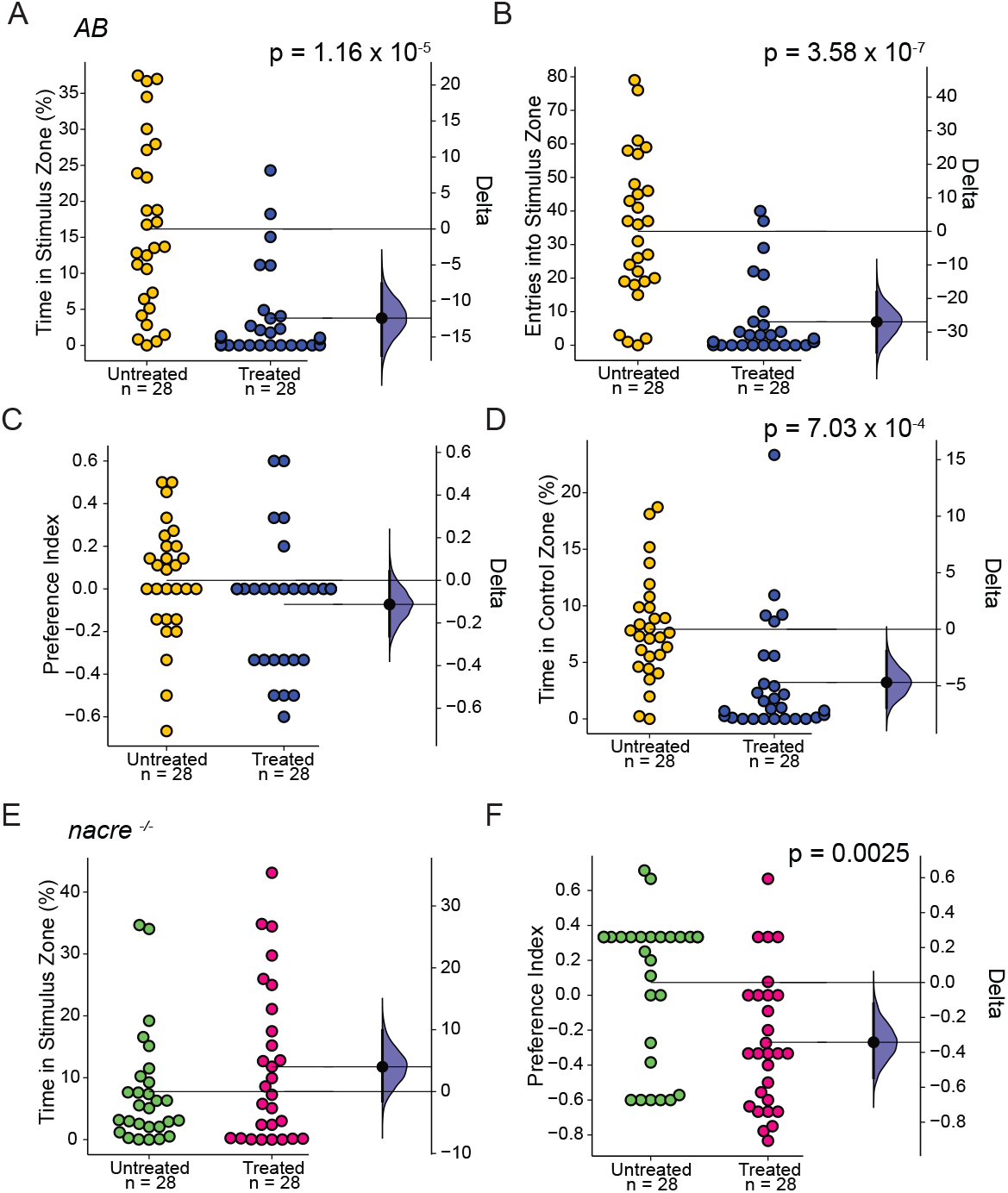
Disulfiram treatment further reduces preference for alcohol. AB WT showed a decrease in (A) time spent in the stimulus zone (delta = −12.3%; 95% CI [−17.6, −7.6]; p < 0.0001; *δ* = −0.7) and (D) control zone (delta = −4.9%; 95% CI [−6.9, −1.9]; p < 0.0001, *δ* = −0.6), and (B) number of entries into stimulus zone, and a moderate decrease in (C) the preference for 5% alcohol after disulfiram treatment (delta = −0.1; 95% CI [−0.3, 0.04]; p = 0.1559; *δ* = −0.3). The nacre-/- mutant showed no change in (E) time spent in the stimulus zone but showed a reduction in preference for (F) 5% alcohol (delta = −0.3 95% CI [−0.5, −0.1]; p < 0.005; *δ* = −0.5). The mean difference is depicted as a black dot, and the 95% confidence intervals are shown as vertical error bars on floating axes. The shaded area shows the bootstrap sampling distribution of the mean difference. Meaningful effects (−0.4 > *δ* > 0.4, with p-values < 0.01) are depicted in the figure.

### Evaluating the role of candidate genes associated with alcohol dependence in human genetic studies

One way to improve our understanding of the neurogenetics of addiction is to perform functional analyses of candidate genes discovered in human genetic studies using animal models. In a pilot study, we identified missense variants predicted to damage protein function in the gene *CCSER1* by Whole Exome Study (WES) of family trios where at least one family member had been diagnosed with a Substance Use Disorder, including Alcohol Use Disorder (AUD; Asharani PV et al., manuscript *in review*). Intronic variants in *CCSER1* have previously been associated with the methadone dose in opioid-dependent individuals and with alcohol dependence in genome-wide gene-by-environment interaction studies of risky behavior [47,48]. Protein sequence variations have also been found in the gene *CCSER1* in selectively bred alcohol non-preferring rats [49]. To better evaluate if zebrafish can be used to study the neurogenetics of addiction with the aid of SAZA, we generated mutants in the *CCSER1* gene using the CRISPR-Cas9 technology. We identified and raised two independent mutant lines, CCSER1 P47F and CCSER1 S48Y*, to adulthood (Figure 6A).

**Figure 6.**
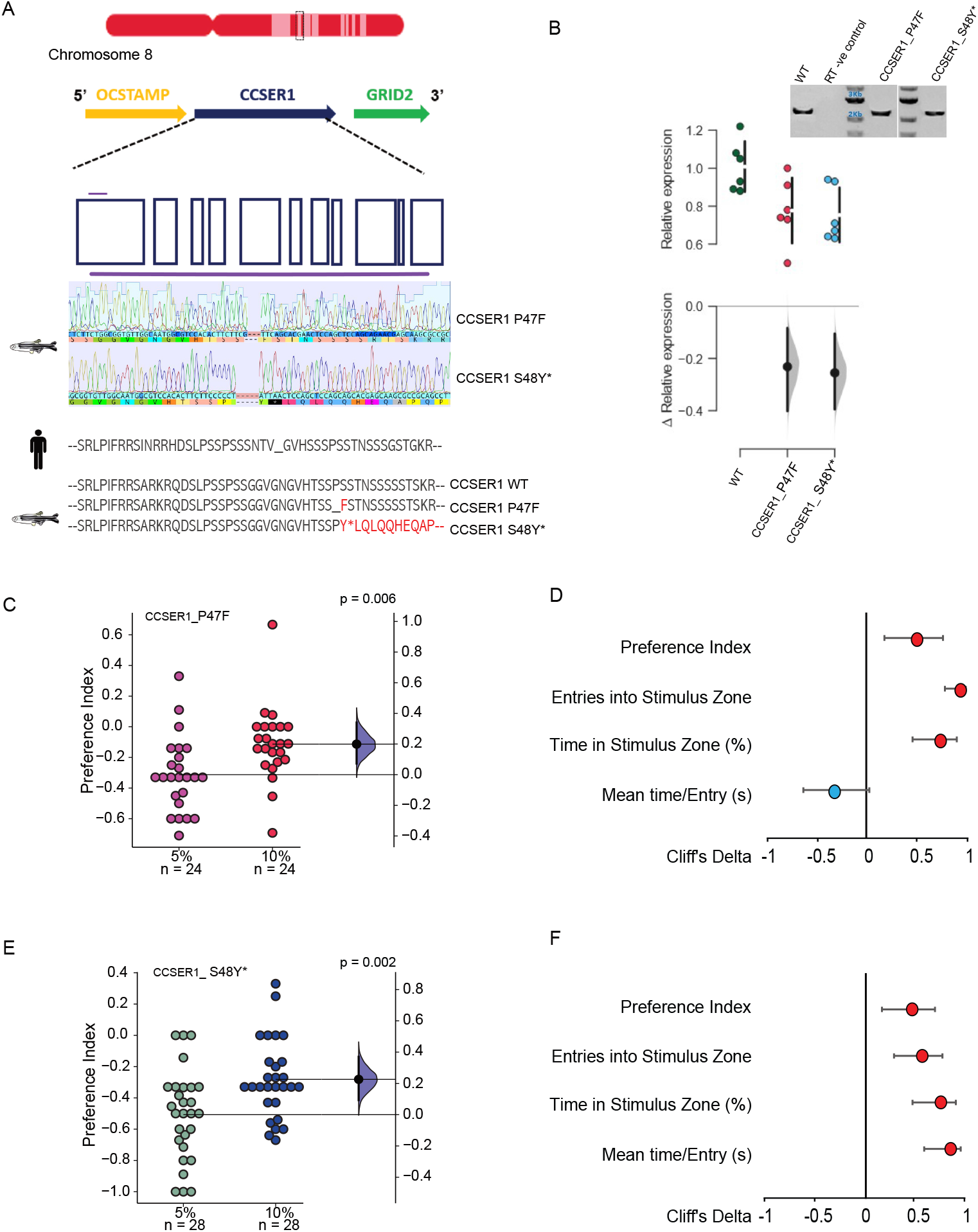
*CCSER1* mutants reduce alcohol self-administration. (A) Schematic shows CRISPR/Cas9 target (in purple) in exon 2 of *CCSER1* gene. Genomic sequence and the protein sequence changed in the two mutant lines generated - CCSER1 P47, and CCSER1 S48Y* is shown below. (B) qRT-PCR showed approximately a 20% reduction in the expression of CCSER1 in mutant brains (n = 6/group; p < 0.05). Inset shows gel pictures of the full-length cDNA from wild-type and the mutants. Both the (C, D) CCSER1 P47, and (E, F) CCSER1 S48Y* mutants show a relative increase in the preference for 10% alcohol compared to 5% alcohol. (C, E) The mean difference is depicted as a black dot, and the 95% confidence intervals are shown as vertical error bars on floating axes. The shaded area shows the bootstrap sampling distribution of the mean difference. Meaningful effects (−0.4 > *δ* > 0.4, with p-values < 0.01) are depicted in the figure. (D) Forest plots for CCSER1 P47 (F) and CCSER1 S48Y* show Cliff’s delta (*δ*) for other parameters. Meaningful effects (−0.4 > *δ* > 0.4, with p-values < 0.01) are shown in red.

The mutant fish were first outcrossed to eliminate potential off-target effects and the heterozygotes in F1 were incrossed to generate F2 homozygous mutants (Figure 6A). Antibodies against zebrafish CCSER1 were unavailable, but quantitative RT-PCR revealed that the *CCSER1* mRNA expression levels were reduced by approximately 20% in both the mutants (Figure 6B). We examined the mutant lines’ responses to both 5% and 10% alcohol in the SAZA and compared them to those of wild-type controls. Both the mutants S48Y* (Supplementary Figure S4A-D), and the P47F (Supplementary Figure S4E-H) showed a reduction in preference of alcohol. This reduction in self-administration preference phenocopies the response of alcohol non-preferring rats that harbor protein function damaging mutations in the *CCSER1* gene [49]. The response of heterozygous and homozygous mutants was comparable (Supplementary Figure S4A, S4E). However, the relative preference for 10% alcohol compared to 5% was increased in both P47F (Figure 6C, mean difference = 0.19, 95% CI [0.07, 0.33], *p* = 0.006), and S48Y* (Figure 6E, mean difference = 0.22, 95% CI [0.08, 0.36], *p* = 0.002) mutant lines. Changes in the other parameters also reflected this increase in the relative preference (Figure 6D, 6F). This is a reversal compared to the wild type response described above. Thus, these results demonstrate that SAZA can be an effective tool to study neurogenetics of addiction and that CCSER1 likely plays a previously unappreciated role in the development of alcohol dependence in humans.

## Discussion

Here, we describe an assay in which the operant behavior of juvenile zebrafish in a two-choice chamber (SAZA) controls their exposure to a stimulus. This assay was used to examine the response of commonly used zebrafish lines to alcohol. Like the alcohol response behavior of many other animals tested in contingent designs, zebrafish showed a biphasic response, preferring exposure to low concentrations and avoiding self-exposure to higher concentrations.

### A two-choice assay system similar to the two-bottle choice assay in rodents

Because of their suitability for use in neurogenetic studies at a young age, zebrafish have been employed in a large number of studies to model the effects of exposure to addictive substances and to characterize withdrawal-like behaviors after chronic exposure to psychoactive substances, including alcohol [17,19,25,50–52]. In these studies effects on locomotion, shoaling, predator avoidance, anxiety-like behaviors, and place preference have been documented. Only two previous studies have observed contingent behaviors to investigate the natural responses of the fish to addictive substances or in the development of dependence. One that utilized an associative training system with an active platform for opioid self-administration is the best example of a contingent assay to date and is comparable to rodent intravenous self-administration experiments [24]. The second measured the consumption of alcohol-containing gelatin pellets [25]. Both of these designs used adults and may also require additional training time.

We have developed an assay that is suitable for fish at a much younger age, and therefore adds a new dimension to addiction studies using zebrafish. The assay is suitable for larvae that have started free-swimming (5 to 7 dpf); however, we have found that, up to 2 weeks of age, fish show higher variability. Therefore, our study used 3- to 4-week-old fish, in which this variability is reduced. The assay delivers a stimulus that is rapidly diluted and cleared from the system unless fish stay in the stimulus zone and continue to trigger the stimulus delivery. The volume of the delivered stimulus is therefore directly proportional to the time spent in the stimulus zone. Our upper estimate suggests there is a 5-10 fold dilution to the stimulus in the assay tank compared with its concentration at the source, although this varies with the molecular weight and specific density of the stimulus (Figure 1). Even though the duration of the assay was short, the effect of alcohol exposure was clear in the post-exposure behavior and quantitative measures during the self-administration phases. We used a preference index (PI) that quantifies the relative preference for the stimulus compared with the control; while other parameters, such as the number of entries into the stimulus zone, percentage time in the stimulus or control zone, mean velocity in the stimulus zone, and the mean time per entry, provide an absolute measurement of the response to each stimulus. Among the designs used in contingent assays, our assay is most comparable to the two-bottle choice assay for rodents [28], in which animals are presented with two bottles of water, one of which is later replaced with a stimulus. In this study, we examined only the acute response of naive animals, but more complex procedures that require longer exposure to the stimulus, multi-day trials, or a schedule of reinforcement can be easily implemented.

### Hormesis response

Our results suggest that the experimental fish regulated their exposure to alcohol to a few seconds at a time, even at the preferred, lower concentrations (approximately 5 seconds at a time; Figure S1). The findings also suggest that, as described previously for other vertebrates and many human behaviors [35], zebrafish show a hormetic response to increasing concentrations of alcohol. The self-administration dose-response curves for alcohol in most animals studied until now have been biphasic, with the effects of alcohol at higher concentrations often showing inhibitory effects [53]. The results presented further suggest that it is not just the effects of alcohol (Supplementary Video 2) that are biphasic, but fish preference for alcohol itself is also biphasic. The ubiquity of finding such non-linear responses in zebrafish behavior [22,34] is interesting to note and relevant for pharmacological studies and suggests that they are likely to be much more universal than previously appreciated [32]. Olfactory stimuli, for instance, are also known to transition from being attractive to aversive depending on their concentration [33,54]. Overall, alcohol is attractive at lower concentrations and, when given a choice, zebrafish avoid exposure to high concentrations.

### Effect of pre-exposure on self-administration

Based on the observations of place preference after a single exposure to alcohol [40,41], we expected pre-exposure to 1% alcohol for 30 minutes for 5 consecutive days to also have an impact. One possibility was an increase in the preference for alcohol in the SAZA. The duration of preexposure was chosen based on previous findings that alcohol levels peak in the body, and particularly the brain, within 20 to 30 minutes after the start of exposure [55,56] and that 5 days of exposure to psychoactive substances often induces notable changes in the physiology and behavior of zebrafish [57]. However, the preference for low-concentration alcohol self-exposure did not change as much as expected for either line (Figure 4). The only notable change was seen in the AB wild-type fish, who displayed an increased frequency of entry to the stimulus zone (Figure 4D, G, H). Nevertheless, as the time spent per entry did not change, the overall effect in terms of the total time in the stimulus zone, or the PI, was unchanged. Hence, we concluded that pre-exposure did not appreciably change the zebrafish’s preference for alcohol exposure.

### Inbred line differences and similarities

A zebrafish line with a *nacre^-/-^* mutation has been widely used in neural activity imaging studies, as the fish lack pigmented melanophores [36–38], and previous studies have shown it to have behavioral differences compared with the AB wild-type [39]. These *nacre^-/-^* mutant fish were generated by N-ethyl-N-nitrosourea (ENU) treatment of AB wild-type fish and are mutant for microphthalmia gene *Mitf* [58]. The *Mitf* gene is known to be a cell fate determinant affecting the pigment cells of the zebrafish neural crest but has not been reported to have a role in alcohol metabolism. Even though alcohol exposure affects the morphology and dispersion of melanocytes [59], we did not expect to see any differences between our inbred lines. However, as our results show, while the overall response was similar, there were distinct quantitative differences (Supplementary Figure S2 compares the two lines directly). The *nacre^-/-^* fish were more tolerant to alcohol and, in general, self-administered higher volumes of all concentrations (directly proportional to the time spent in the stimulus zone) with or without prior experience of alcohol. This was also reflected in only a modest reduction in preference after disulfiram treatment of *nacre^-/-^*, while the preference change was much more pronounced in AB fish. The exact mechanisms behind why *nacre^-/-^* mutants differ are unclear at present. One possibility is that these differences are consequences of *nacre^-/-^* mutants having been maintained as an inbred line for many tens of generations. Previous studies have noted substantial quantitative differences among inbred lines, maintained as short fin, long fin, or leopard strains [60], and the influence of the genetic background when comparing the effects of alcohol [61]. Similar strain differences have also been noted in rodents [62,63]. The second possibility is that the *nacre^-/-^* mutation has a direct role. The interplay between alcohol and cAMP signaling [64] has been implicated in triggering aggregation [65], and this interplay may also regulate alcohol sensitivity or tolerance in the body [66,67]. Future studies examining the blood alcohol concentrations and melanocyte-stimulating hormone levels of the zebrafish line will be highly valuable in teasing apart these differences. Nonetheless, it suggests that the assay is sufficiently sensitive to evaluate subtle differences between animals from different genetic backgrounds.

### Functional CCSER1 is necessary for normal response to alcohol

Our results suggest that mutations in the gene *CCSER1* (HUGO gene ID HGNC:29349) have an impact in the development of AUD. The gene previously known as *FAM190A* has been studied extensively for its role in cell division and tumorigenesis [68–70]. As the intronic variants in this gene were associated with multiple Substance Use Disorders (SUDs) and risky behavior [47,48], and the disruptive exonic variants were discovered in WES study (Asharani PV et al., manuscript *in review*), it made for a good candidate for examination in a model system. Linkage analysis of selectively bred rats had identified potentially damaging missense mutation in the *CCSER1* gene (P82L) in alcohol non-preferring rats in a previous study. This study reported that preference for alcohol was reduced by approximately 25% when rats carried mutant copies of *CCSER1* (or *FAM190*) and 4 other genes. Mutant fish in our study also showed a reduction in preference for administering alcohol. Thus, our results replicate the earlier findings. This decrease in preference in the mutant fish is limited to alcohol and is not observable for other addictive substances such as nicotine (data not shown). In the human WES study, however, protein function damaging mutations were associated with alcoholism. Our final set of results of the reversal in the relative preference for higher concentrations when compared to the wild-type response suggests a possible explanation. A similarity in its role in 3 distant species is remarkable. Further studies are needed to unravel the molecular mechanisms connecting CCSER1 protein function and alcohol preference.

One limitation of our assay system is that it cannot estimate the amount of alcohol (or a drug) *in vivo* in the animal, as it does not involve intravenous injections; however, it is also an advantage, as any water-soluble drug is introduced non-invasively. Another limitation is that, although alcohol delivery is contingent, the assay does not directly evaluate a “drug-seeking” behavior in this experimental design. Further modifications to the design, such as a chronic multi-day testing regimen along with a progressive ratio schedule, will be required to address such issues.

Even with these limitations, the use of a two-choice contingent assay can be employed for quantitative neurogenetics studies of addiction and for screening novel small molecule deterrents and cessation aids. SAZA adds to the growing arsenal of zebrafish-based tools that can be applied in personalized medicine research. Our results and the SAZA expand the utility of zebrafish for examining the consequences of candidate gene mutations associated with the development of addiction that have been identified in human genetic studies. Any changes in the natural responses of fish limiting their self-exposure or the attractiveness of low concentrations will be easy to identify. Similarly, the effectiveness of a small water-soluble molecule postulated to act as a deterrent can also be rapidly screened before it is tested in other preclinical models, reducing both the cost and duration of testing.

## Materials and Methods

### Fish husbandry

A total of 392 *nacre^-/-^* fish and 224 AB wild-type fish between the ages of 3 to 4 weeks were used in this experiment. The fish were bred and housed in the laboratory fish facility (Institute of Molecular and Cell Biology, A*STAR) in groups of 20 to 30 in 3-L tanks under standard facility conditions. All experiments were performed following guidelines recommended by the Institutional Animal Care and Use Committee (IACUC) of the Biological Resource Center at A*STAR. Approved experimental protocols (IACUC 171218; 201529) were followed.

### SAZA experimental setup

Videos were acquired at 30 fps on an acA2040-90μM USB3.0 : Basler Camera for the two-choice SAZA. The entire setup was backlit with a white LED lightbox (LightPad LX Series, Artograph, USA). The assay chamber shown in Figure 1A, which was designed in-house, had internal dimensions of 76 × 32 × 30 mm (L × W × H). This gave about 7 body-lengths in length and 4 in width. The tank was fabricated from 3-mm thick opaque acrylic sheets that reduced the internal reflective surface when filled with water. Two choice zones were created using a partition (30 × 2 × 25 mm [L × W × H]) at one end of the chamber (Figure 1A; green boxes). The delivery of the stimulus or control (alcohol or system water, respectively) was controlled by solenoid pinch valves (Automate Scientific, USA, SKU: 02-pp-04i) with silicon tubes (sold as tubing with 1/16” outer diameter and 1/32” inner diameter) connected to 10-mL syringe reservoirs. A suction tube connected to the partition in the middle of the chamber served as an outflow. This arrangement created three virtual zones that were chemically separated but permitted unhindered physical swimming access (Supplementary Video 1). Fish were tracked online using custom-written LABVIEW software CRITTA (http://www.critta.org) as previously described [33]. For the experiments described here, CRITTA was configured to create a closed-loop system, such that the entry of the fish into the virtual zones triggered the pinch valve delivery system. As the outflow suction constantly extracted the liquid, the delivered stimulus was rapidly diluted (Figure 1E) unless the fish re-entered a virtual zone.

### Assay design

The assay chamber was filled with 40 ml of system water at the start of the experiment. Tubes delivering control system water or the stimulus were placed at one end of the assay tank (Figure 1A; black arrows). To eliminate bias due to the stimulus delivery location, the stimulus delivery zone was randomly assigned to either the left or right of the assay chamber for half of the subjects. Once the video recording setup was ready, pairs of fish were collected from their home tanks and gently delivered individually into assay chambers. Each trial lasted 24 minutes (Figure 1B, C) divided into three phases – pre-exposure, self-administration stimulus delivery, and post-exposure phase. During the stimulus delivery period, entry of the fish into either the virtual stimulus zone or control zone triggered the delivery (Figure 1B; Supplementary Video 1), thus fish could freely choose to self-administer either the stimulus or control solutions. The apparatus was washed thoroughly after each set of fish was tested. A total of 28 fish subjects naive to the assay system were tested per condition.

### Analysis

Custom scripts written in Python were used to analyze data in a semi-automatic manner. The analysis scripts also generated graphics and spreadsheets with data on time spent in each of the virtual zones, velocity, number of entries, and percentage time spent. The PI was also calculated based on the volume of the stimulus delivered as [(volume of stimulus delivered - the volume of control delivered)/(volume of stimulus delivered + volume of control delivered)]. A PI of + 1 thus indicated a maximum relative preference for the stimulus, while a PI of −1 indicated the maximum relative avoidance of the stimulus. Analysis script output included sparklines to represent the number and duration of entries into the stimulus or control zone (Figure 1B).

### Stimulus dilution in the assay chamber

The extent to which the three virtual zones of the assay chamber remained chemically distinct was examined through the course of the experiment by using solutions with colored dyes as stimuli. This approach permitted an estimation of the dilution of the stimuli in the assay chamber during the self-administration phase. Three different dyes with molecular weights of 604.46 Da (Artificial Cochineal Red), 576.62 Da (Artificial Apple Green Colour), and 477.38 Da (Artificial Egg Yellow Colour) were used. Based on the measurable absorbance at 595 nm of a serial dilution of green food dye (576.62 Da; dilutions from 0.1 to 2 × 10^−5^ % w/v in system water), 0.01% w/v in 1% alcohol was used as the solvent for the dyes. Water samples from four regions (Figure 1D) of the chamber were collected in duplicate every 3 minutes over the 18-minute self-administration phase (Figure 1C). Representative absorbances recorded during one such test run are shown (Figure 1E). This experiment revealed that after stimuli were delivered into the stimulus zone (position 1 in Figure 1D) they were diluted by 5-10 fold compared to the concentration at the source. The stimulus was rapidly diluted to a fraction (Figure 1E) outside this delivery zone (positions 3 and 4 in Figure 1D). Individual fish sometimes favored either the left or right delivery zones, even when both zones delivered system water; therefore, all experiments used a balanced design, in which the stimulus delivery zones were switched between the two zones for half of the fish. In all cases, fish entry into the non-stimulus zone triggered the delivery of control or system water. The volume of stimulus delivered was linearly proportional to the duration of valve opening (Figure 1F), which in turn, was determined by the duration the fish subjects remained in the stimulus or control zones.

### Pre-treatment before the acute test in SAZA

In the alcohol pre-exposure experiments, 3-week-old fish were placed in holding tanks in 500 ml of system water in groups of 28 and exposed to either 1% alcohol or system water for 30 minutes every day for 5 days. Fish were tested with the SAZA on day 8. The pre-treatment alcohol concentration was based on a description of suitable concentrations for obtaining quantifiable effects within 20 minutes [59].

For disulfiram (Antabuse, Sigma, PHR1690) pre-treatment, 3- to 4-week-old fish were exposed to either 500 nM disulfiram or system water overnight. Twenty-eight fish were placed in static tanks with approximately 500 ml of system water for both the control and treatment groups. Disulfiram was prepared by making a 10-μM stock solution, which was diluted in system water to obtain a final concentration of 500 nM.

### Statistical analysis and interpretation of behavioral data

The use of null hypothesis significance testing and an over-reliance on *p*-value-based dichotomous thinking for rejecting or accepting hypothesis has been extensively criticized [71,72]. An alternative recommendation is to measure and report the difference between two groups in the form of the effect size [72]. To interpret the results, traditional significance testing was supplemented with estimation statistics and Gardner-Altman plots to determine the effect sizes and accuracy of the measurements, as suggested by a recent report [72]. For readers unfamiliar with these plots, the primary (left) axis represents the measured parameter for both groups, and the distribution of individual data points is shown as swarm plots. The floating difference-axis, either on the right or below, shows the delta values of the groups being compared. A black circle represents the mean of the delta, the whiskers represent the 95% confidence intervals, and the shaded area shows bias-corrected and accelerated 5,000 bootstrap samples confidence intervals. Adobe Illustrator (Illustrator, 2018) was used to change plot colors and font. Permutation *t*-tests are reported. Additionally, forest plots showing Cliff’s delta were used to visualize and compare multiple parameters [73]. `Overall, a Cliff’s delta effect size greater than 0.4 (or less than −0.4) and a` *p*-value of < 0.01 were used in this study to identify meaningfully large and practically relevant results, as suggested by a previous study (Goodman et al., 2019; Halsey, 2019). A *p*-value of < 0.01 coupled with smaller effects of 0.2 to 0.4 for Cliff’s Delta, which were also considered potentially valuable but provisional until more experiments are conducted, are shown in purple. Results are reported as delta = XYZ, 95% CI [lower, upper], and *p*-values, and Cliff’s delta when reported as *δ*.

### CRISPR/Cas-9 based mutagenesis to generate CCSER1

A guide RNA (gRNA) against exon 2 of CCSER1 with the following targets GGAGTTCGTGCTGGAGGGGG, or ATTGCCAACACCGCCAGAAG were used to generate indel mutations. gRNA template was synthesized using PCR and transcribed with MEGAscript T7 transcritption Kit (Life Technologies, AM1344) according to the manufacturer’s protocol and purified by ammonium acetate precipitation. The Cas9 mRNA was made by linearizing the expression vector pT3TS-nCas9n (Addgene#46757) with XbaI restriction enzyme (NEB) and invitro transcription using T3 Message Machine transcription (Ambion). Cas9 mRNA was then capped and polyadenylated to protect the mRNA molecule from enzymatic degradation and aid transcription termination. Lithium chloride precipitation was utilized to remove access NTPs from Cas9 mRNA at −20 °C overnight. One microgram of linearized plasmid template typically yielded about 1 - 1.3 *μ*g/*μ*L of Capped mRNA. Cas9 mRNA was resuspended in 30 *μ*L of nuclease free water and stored similarly to sgRNA. 1 nL of the mixture containing 1 μl of gRNA (4 - 5 μg) and 1 μl of Cas9 mRNA (1 μg) was injected into the yolk of 1-cell *nacre^-/-^* zebrafish embryos.

Two independent lines were selected, raised to adulthood and outbred to wild type fish to remove off-target effects if any. F1 adults were incrossed to generate mutant F2. Behavioral experiments were performed on F2 or F3 larvae. Homozygous F2 mutant fish did not survive a reproductive age, but all fish were genotyped after behavioral analysis. No difference in the homozygous and heterozygous animal behavioral responses were detectable (Supplementary Figure S4).

### RNA analysis

2-month old zebrafish brains dissected in PBS pH 7.0 were used for RNA extraction. 6 zebrafish per group (wild-type, CCSER1 P47F, and CCSER1 S48Y*) were analyzed. Tissue was homogenized in lysis buffer consisting of Trizol and RNA carrier. RNA was extracted with PureLink Micro Kit (ThermoFisher) based on the manufacturer’s protocol. The concentration and purity of the RNA was determined using NanoDrop 2000 Spectrophotometer (ThermoFisher). Each brain sample yielded approximately 100 ng/μL of RNA. *Quantitative real-time - PCR* (*qRT-PCR*)

The relative expression of the CCSER1 was examined by qRT-PCR. cDNA was prepared by SuperScript II First-strand Synthesis System (Invitrogen). The cDNA was diluted with nuclease-free water to 100 ng/μL. The qRT-PCR amplification mixture (20 μL) contained 100 ng of cDNA, 10 μL 2X Go TaqqPCR Master Mix (Promega), and 300 nM forward and reverse primer. Reactions were carried out in triplicates using 7500 Real-time PCR System (Applied Biosystems) in a 96-well plate. The data were averaged and normalized to β-actin and Elif to obtain the ΔCT values (geomean). All PCR efficiencies were above 95%. Sequence Detection Software (version 1.3; Applied Biosystems) results were exported for calculations.

## Acknowledgments

We would like to thank the zebrafish fish facility staff at the IMCB, A*STAR for fish husbandry. We also thank Maleyka Mammadova for generating customized analysis scripts.

## Author contributions

Conceptualization: ASM;

Methodology: ACC, JS, ASM

Investigation: FMN, CK, TG

Visualization: ASM, FMN

Supervision: ASM

Writing—original draft: ASM

Writing—review & editing: ACC, JS, FMN, CK, ASM

## Data and materials availability

All data are available in the main text or supplementary materials.

## Supplementary Materials

Supplementary data includes 7 items (4 figures, 1 table, and 2 movies).

