## Supplemental Material for "Contingent-behavior assay to study the neurogenetics of addiction shows zebrafish preference for alcohol is biphasic"

**This PDF file includes:**

Supplementary Text

Figures S1 to S4

Tables S1

Movies S1 to S2

**Other Supplementary Materials for this manuscript include the following:**

Movies S1 to S2

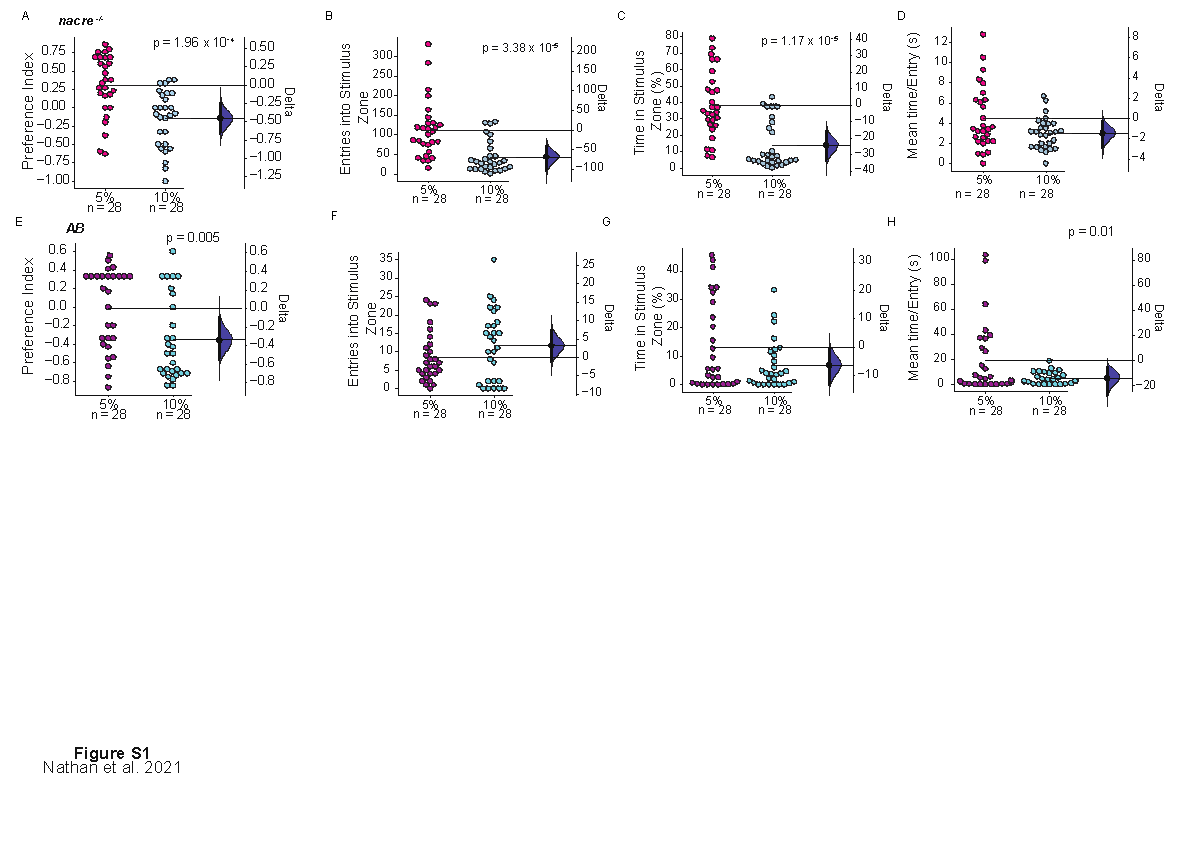
**Figure S1. Zebrafish show a decrease in preference for higher alcohol concentrations**. Data shown in Forest Plots in Figure 3 (in the main text).  (A-D) *nacre^-/-^* and (E-H) AB WT strains showed a preference for 5% alcohol and avoided 10% alcohol. *nacre^-/-^* showed a decrease in the (A) preference index, (B) the number of entries, (B) the percentage time, and (C) the average time per entry, into the stimulus zone when 10% alcohol was used as stimulus in comparison to 5% alcohol. AB WT strain also showed a decrease in (E) preference index, and (H) mean time per entry while (F) the entries increased (G) percentage time, into stimulus zone did not show large changes. In all panels, the mean difference is depicted as a black dot, and the 95% confidence intervals are shown as vertical error bars. The shaded area shows the bootstrap sampling distribution of the mean difference. Meaningful effects (−0.4 > 𝛿 > 0.4, with *P*-values < 0.01) are depicted in the figure.

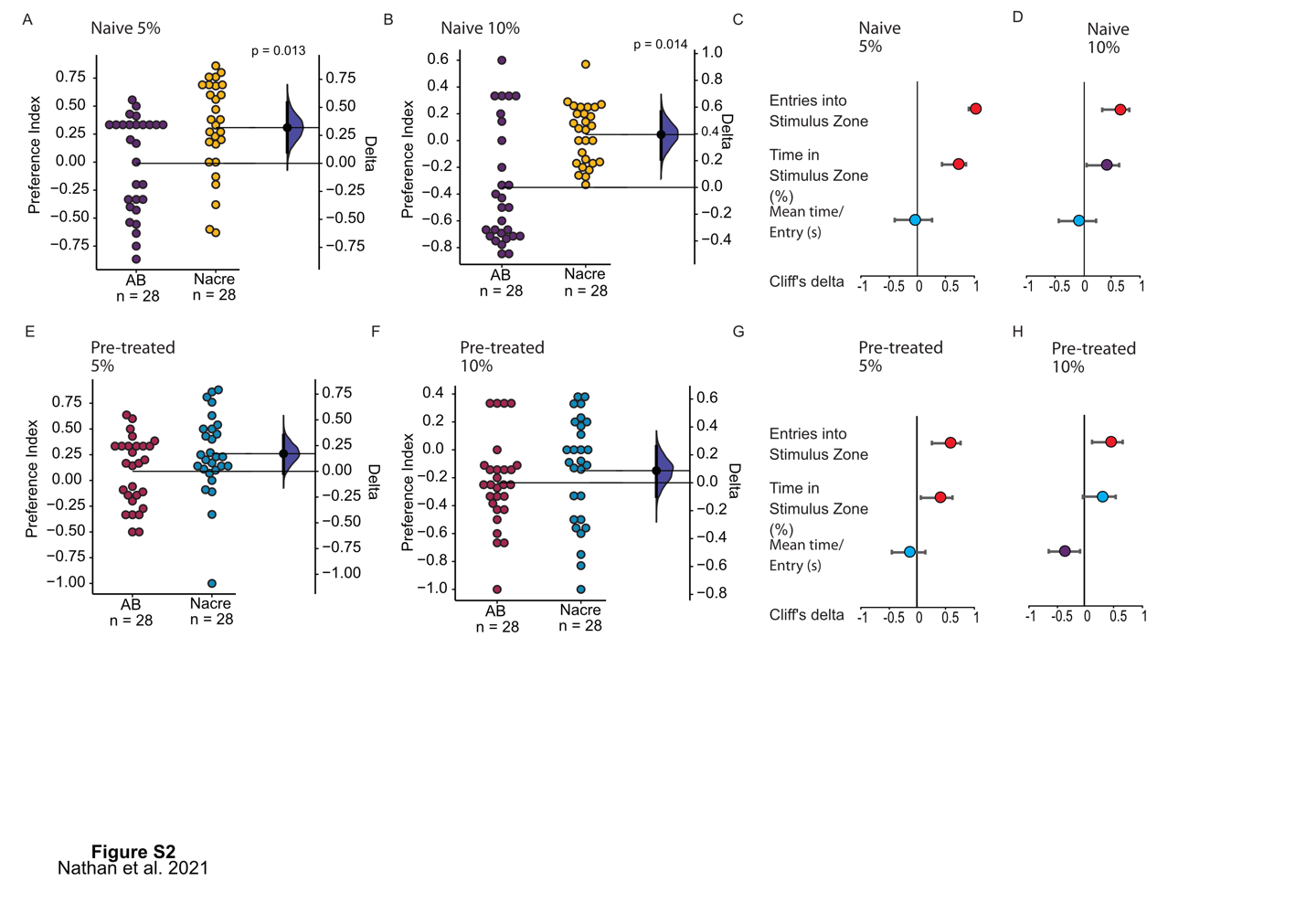

**Figure S2. Genetic background affects absolute volumes and preference for alcohol**. *nacre^-/-^* fish showed greater preference for (A) 5%, and (B) 10% alcohol compared to AB WT when naïve to alcohol. This difference reduced when fish are pre-exposed to alcohol for both (E) 5% and (F) 10% alcohol. Forest plots show Cliff’s delta (𝛿) for other parameters measured for (C) 5% alcohol, and (D) 10% alcohol when the two lines without pre-exposure to alcohol. Forest plots show Cliff’s delta (𝛿) for other parameters measured for (G) 5% alcohol, and (H) 10% alcohol when the two lines were pre-exposed to alcohol. The mean difference in (A, B, E, F) is depicted as a black dot, and the 95% confidence intervals are shown as vertical error bars. The shaded area shows the bootstrap sampling distribution of the mean difference. Meaningfully large effects in (C, D, G, H) are shown in red (−0.4 > 𝛿 > 0.4, with *P*-values < 0.01). Moderate effects tentatively accepted in purple and blue shows insignificant changes.

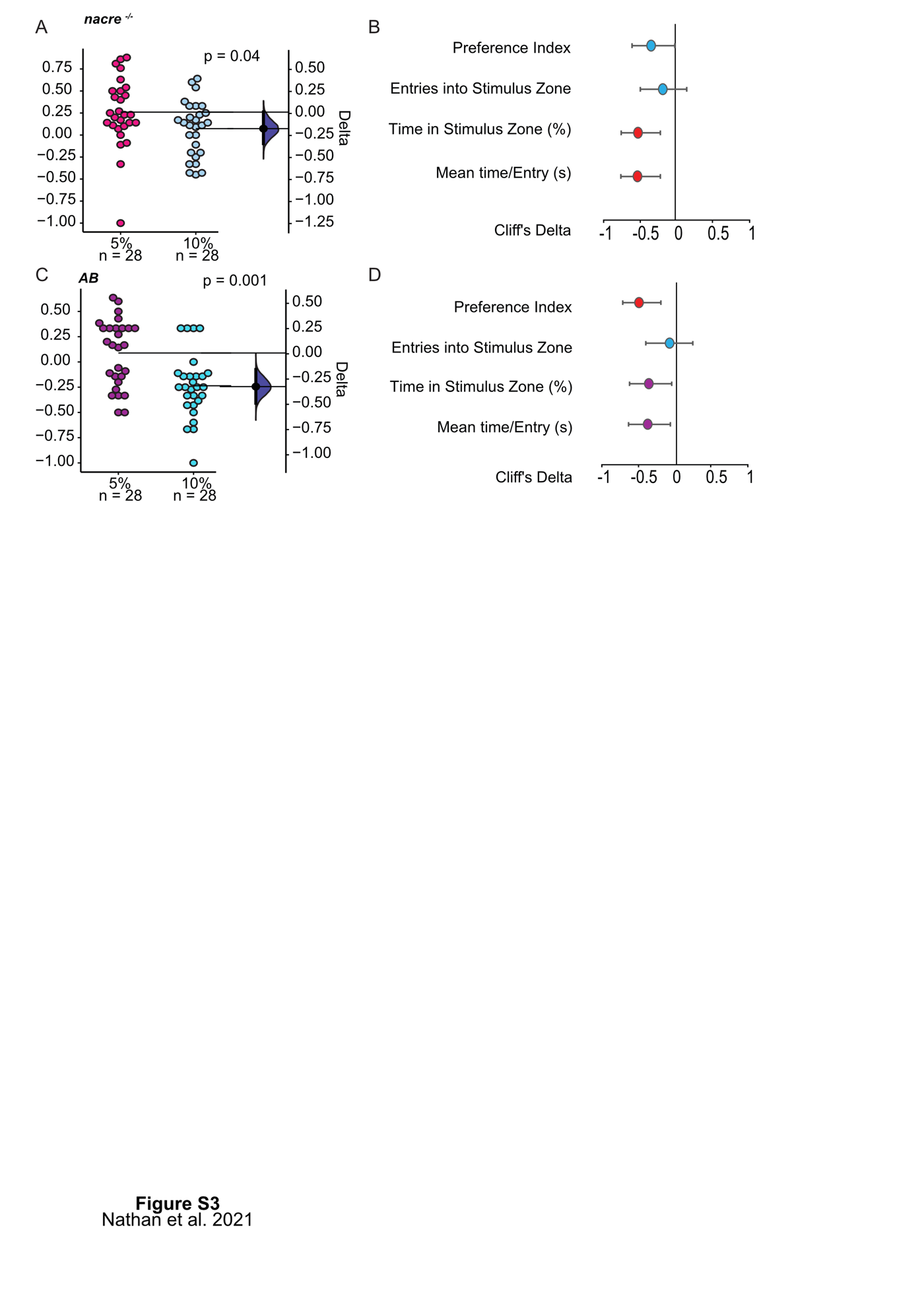

**Figure S3. Pre-exposure to alcohol does not change the preference for low concentration of alcohol.** PI delta for (A) *nacre^-/-^* (delta = −0.2; 95% CI [-0.4, 0.0]; 𝛿 = −0.3) and (C) AB WT = -delta = −0.3, 95% CI [-0.5 -0.1]; 𝛿 = -0.5), and Forest plots for (B) *nacre^-/-^*  and (D) AB WT shows that both inbred line fish continue to prefer 5% alcohol even after pre-exposure to 1% alcohol for 5 days. *nacre^-/-^*  fish spent more time in stimulus zone, but their preference for 5% alcohol was reduced. The mean difference in (A, C) is depicted as a black dot, and the 95% confidence intervals are shown as vertical error bars. The shaded area shows the bootstrap sampling distribution of the mean difference. Forest plots for (B) *nacre^-/-^* and (D) AB WT show Cliff’s delta (𝛿) for other parameters. Meaningfully large effects in (C, D, G, H) are shown in red (−0.4 > 𝛿 > 0.4, with *P*-values < 0.01). Moderate effects tentatively accepted in purple and blue shows insignificant changes.

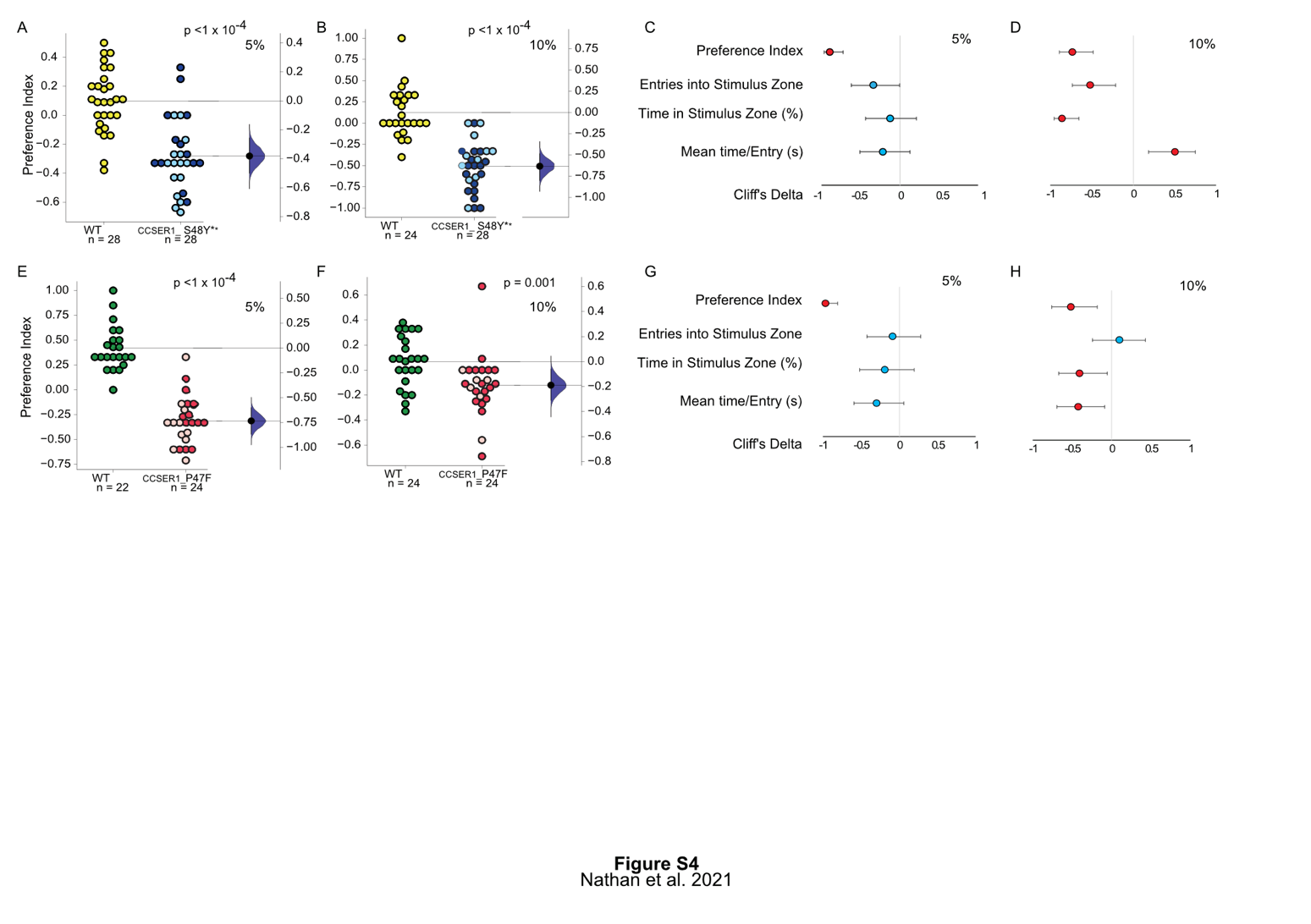

**Figure S4. *CCSER1* mutants have a reduced preference for alcohol.** Both CCSER1 S48Y* and CCSER1 P47F show a reduced preference for (A) 5% alcohol (delta = -0.38; 95% CI [ -0.50, -0.26]; p , 0.001); (B) 10% alcohol (delta = -0.63; 95% CI [ -0.79, -0.49]; p < 0.001), and (C) 5% alcohol (delta = -0.73 95% CI [ -0.86, -0.59]; p < 0.001) and (B) 10% alcohol (delta = -0.28; 95% CI [ -0.43, -0.11]; p = 0.001) respectively compared to wild-type siblings. Forest plots for (C) S48Y* and (G) P47F show that both lines show only a minor change in other parameters compared to wild-type sibling response to administering 5% alcohol. Forest plots for both (D) S48Y* and (G) P47F show that moderate, but significant reduction in the time spent in the stimulus zone when administering 10% alcohol compared to wild-type siblings. (A, B, E, F) Homozygous fish identified by genotyping after the experiment are shown in lighter shade. The mean difference is depicted as a black dot, and the 95% confidence intervals are shown as vertical error bars. The shaded area shows the bootstrap sampling distribution of the mean difference. Forest plots for (C, D, G and H) show Cliff’s delta (𝛿) for other parameters. Meaningfully large effects in (C, D, G, H) are shown in red (−0.4 > 𝛿 > 0.4, with *P*-values < 0.01). Moderate effects tentatively accepted in purple and blue show insignificant changes.

**Table S1.**

Effect sizes and *P*-values for Figure 2 with a shared control (0% alcohol) and five-point dose-response curve (2.5%, 5%, 10%, 40%, 70% alcohol).

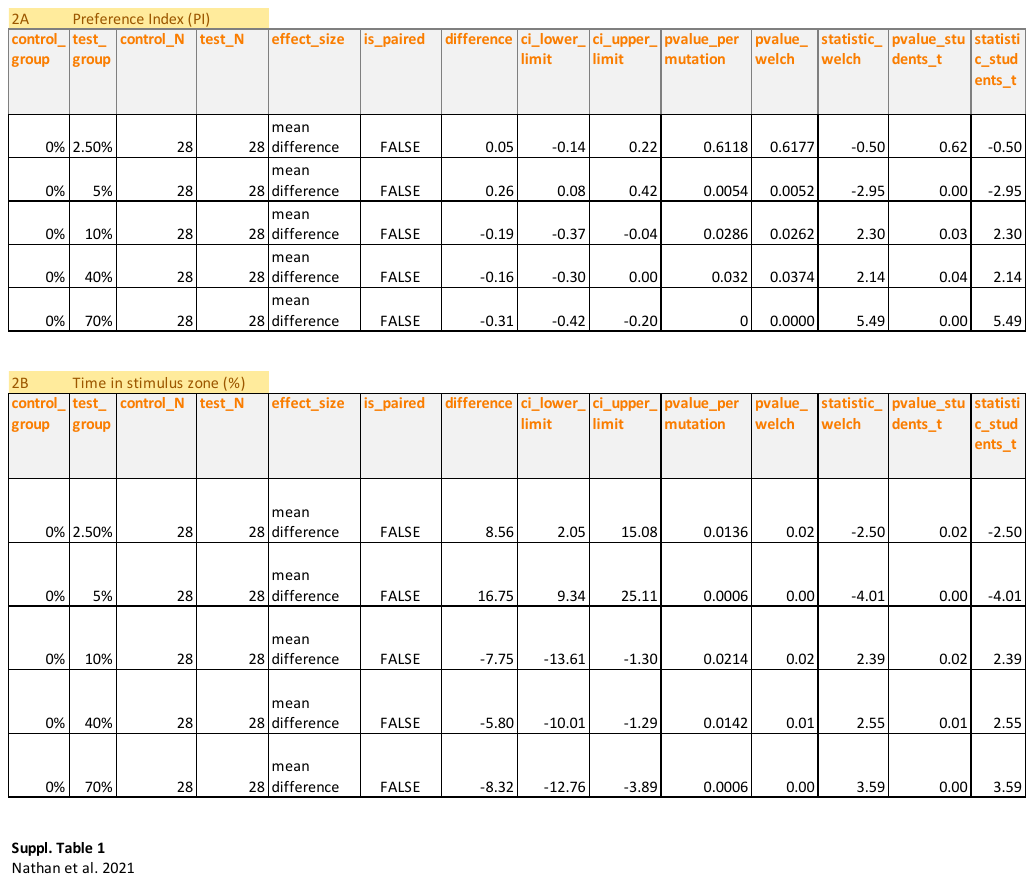

**Movie S1. Two-Choice Self-Administration Zebrafish Assay (SAZA) behavioural setup.** The videos shows the setup that has 3 virtual zones: a stimulus zone, a control zone, and a neutral zone. 3- to 4-week old zebrafish swam freely in the SAZA and triggered the delivery of stimulus or system water from tubes in the delivery zones. Fresh system water was delivered at a constant rate of 1 ml/min from one end, and water was extracted in the middle.

The video uses a colored dye to illustrate the stimulus delivery triggered by the subject fish’s entry into the stimulus zone and its rapid removal from the extraction port.

**Movie S2. Biphasic effect of 1% alcohol on juvenile zebrafish.** Increased shoal cohesion, and coordinated swimming patterns observed by 5-10 minutes of exposure to alcohol. Fish transition to disoriented swimming over time with little social cohesion, observable by 20- 30 minutes.
